# Spatio-Temporal Control of GFP-tagged Protein Degradation in the Mouse

**DOI:** 10.1101/2025.04.17.649420

**Authors:** Alexandra Prado-Mantilla, Terry Lechler

## Abstract

Loss of function studies are a central approach to understanding gene/protein function. In mice, this often relies upon heritable recombination at the DNA level. This approach is slow and non-reversible, which limits both spatial and temporal resolution of analysis. Recently, degron techniques that directly target proteins for degradation have been successfully used to quickly and reversibly knockdown proteins. Currently, these systems have been limited by lack of tissue/cell type specificity. Here, we generated mice that allow spatial and temporal control of GFP-tagged protein degradation. This Degron^GFP^ line leads to degradation of GFP-tagged proteins in different cellular compartments and in distinct cell types. Further, it is rapid and reversible. We used Degron^GFP^ to probe the function of the glucocorticoid receptor in the epidermis and demonstrate that it has distinct functions in proliferative and differentiated cells – an analysis that would not have been possible with traditional recombination approaches. We propose that the ability to use GFP knock-in lines for loss of function analysis will provide additional motivation for generation of these useful tools.

## Introduction

Loss of function approaches are important for determining the roles that genes and their products play. In mice, genetic loss of function is commonly induced by recombination using Cre/Lox or FLP/FRT technology. The advantage of these approaches is that they allow both cell-type control through tissue-specific promoters and can offer temporal control through pharmacological induction of recombination. Recombination results in a heritable alteration in the genome and is therefore passed down to progeny cells in a non-reversible manner.

There are also drawbacks to recombination-based gene disruption. Since this method operates at the genome level, the preexisting mRNA and protein products remain unaffected at the time of recombination. Thus, the half-lives of these products may result in a slower protein depletion rate compared to the turnover of certain tissues, significantly hindering loss of function analysis. For example, villar epithelial cells of the intestine are replaced every 2-3 days, shorter than the half-life of many proteins (Kopf & Sixt, 2019). To overcome this problem, recombination is often induced in stem/progenitor cells resulting in mutant progeny. This raises additional caveats, as it becomes difficult to identify the cell type(s) where gene function is required.

Differentiated cells can regulate the cell behavior of their progenitors (Ning et al., 2021), therefore, determining whether genes are acting cell autonomously in stem cells or non-autonomously through their differentiated progeny is challenging. Further, recombination is not reversible. This can make it difficult to identify time intervals when gene function is required, and restoring gene function involves additional alleles.

These problems can be circumvented by directly targeting proteins using tags that promote their degradation: degron tags (Harris & Trader, 2025). Numerous *in vitro* studies have demonstrated that attaching degron tags to a protein of interest leads to their rapid proteasomal degradation (Macartney et al., 2017; Natsume & Kanemaki, 2017). Degron-fused proteins can be conditionally degraded in the presence of specific ligands, such as Auxin-Inducible Degradation (AID) and dTAG technologies, providing temporal control of degradation (Nabet et al., 2018; Nishimura et al., 2009). In the last few years, mouse models that conditionally induce acute protein depletion using AID (Macdonald et al., 2022; Yesbolatova et al., 2020) (Suski et al., 2022) and dTAG technologies (Abuhashem et al., 2022) (Bisia et al., 2023; Bondeson et al., 2022; Nabet et al., 2018; Olsen et al., 2022) (Hernandez-Moran et al., 2024) have been generated, demonstrating the utility of this approach. These methods were both rapid and reversible, in contrast to recombination. However, they did not allow tissue-specific targeting. Furthermore, these approaches require the creation of new mouse lines with specific degron tags for each gene of interest, which can be time-consuming and costly. Thus, mouse tools that allow more general targeting, especially with a tag that provides additional functionality, are needed.

A number of GFP knock-in lines already exist and allow analysis of expression, protein localization, and dynamics. We sought to address whether they could also be used for loss of function analysis. Here, we have generated mouse lines for targeted degradation of GFP-tagged proteins. By making this system antibiotic-inducible and under the control of a tissue specific promoter, we are able to rapidly induce protein degradation with temporal and spatial specificity, as well as reversibility. Employing this tool, we were able to degrade endogenous GFP-tagged proteins in different tissues and cell compartments in mice. We show the utility of this approach by knocking down the glucocorticoid receptor (GR) in either proliferative or differentiated cells within the epidermis to compare its phenotypes to published GR epidermal knockouts. We demonstrate that GR has distinct functions each of these cellular compartments. Due to the time and costs associated with generating knock-in/knock-out mouse lines, the ability to generate one strain that allows both imaging and loss of function abilities will be advantageous.

## Results

### Degron^GFP^ degrades GFP-tagged protein in different cellular compartments

To target GFP-tagged proteins, we took advantage of the AdPROM system which contains a cassette encoding the substrate recognition element of E3 complex VHL, fused to an anti-GFP nanobody (VHL-αGFP) under the tetracycline-responsive promoter element (TRE) (Fulcher et al., 2016; Macartney et al., 2017). The GFP nanobody binds to a GFP-tagged protein, bringing it into close proximity to the E3 ligase for its polyubiquitination and subsequent degradation (Figure 1A). When used in combination with tetracyline/doxycycline inducible lines (Tet-ON), tissue specificity can be achieved by directly controlling the expression of the Tet-regulated transactivator (rtTA) under a tissue-specific promoter (Urlinger et al., 2000), or by crossing it to a tissue-specific Cre line along with a floxed ROSA-rtTA line (Hochedlinger et al., 2005). Therefore, this system is expected to allow both spatial and temporal control of protein degradation in mice. We generated mouse lines expressing this construct, hereafter referred to as Degron^GFP^.

**Figure 1.**
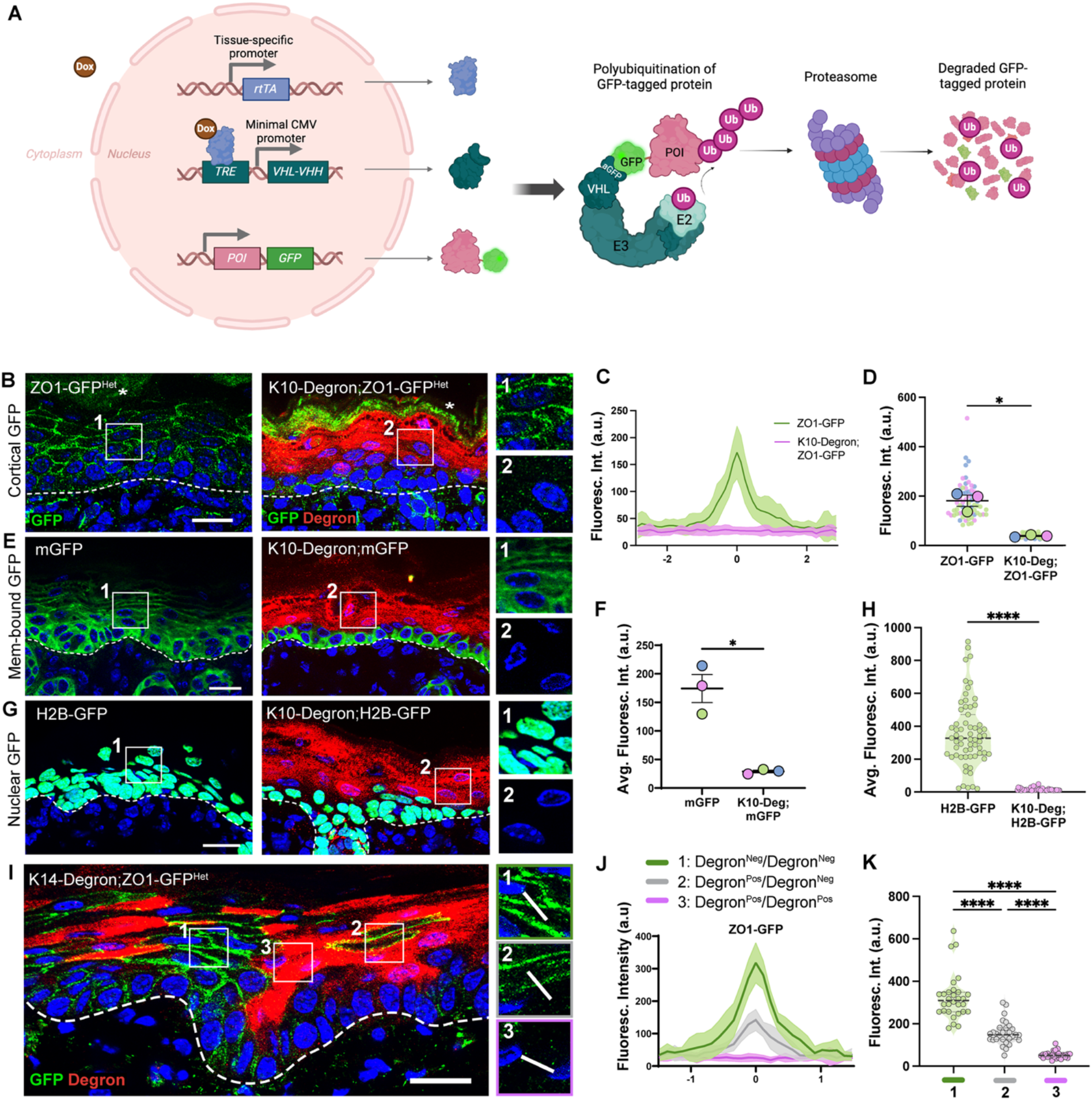
Degron^GFP^ degrades GFP-tagged proteins located in different cell compartments: (A) Diagram of the Degron^GFP^ system. To the left, alleles required for the system: rtTA under a tissue-specific promoter, TRE-VHL-aGFP, and gene of interest fused to GFP (for loss of function studies, two alleles of GFP knock-in are required). To the left: Degron’s mechanism of action; briefly: VHL-aGFP (degron), a component of the E3 ligase complex, is translated only upon Dox exposure and binds to the GFP-tagged protein of interest (POI), bringing it to proximity to E2 to facilitate the transfer of ubiquitin. Polyubiquitinated POI-GFP is then sent for proteasomal degradation. (B) Immunofluorescence staining of GFP (green) and degron (red) in P0 back skin epidermis of K10-Degron;ZO1-GFP neonates and control ZO1-GFP littermate. Insets show only GFP in control (1) or in K10-Degron;ZO1-GFP (2). Dotted lines represent the basement membrane. *: Indicates auto-fluorescent cornified layer. All scales: 20μm. (C) GFP fluorescence intensity across cell-cell junctions in the epidermal suprabasal layers of ZO1-GFP (green), and suprabasal degron^Pos^ areas of K10-Degron;ZO1-GFP neonates (pink). Line scan graph shows measurements across 12 cell-cell boundaries from 3 different mice per genotype. Data are represented as mean±SD. (D) Quantification of fluorescence intensity at suprabasal-suprabasal cell boundaries in ZO1-GFP, and degron^Pos^ areas in K10-Degron;ZO1-GFP neonates. Each dot represents the maximum intensity at cell-cell junctions, large circles represent the average for each mouse (n=3 mice per genotype, at least 20 line scans per animal). Lines show the mean±SEM. p=0.0294, paired two-tailed t-test. (E) Immunofluorescence staining of GFP and degron in P0 backskin epidermis of K10-Degron;mGFP neonates and control mGFP littermate. Insets show only GFP in control (1) or in K10-Degron;mGFP (2). (F) Quantification of average mGFP fluorescence intensity per mouse from all suprabasal areas in controls (n=3 mice) or degron+ suprabasal areas in experimental neonates (n=3 mice). Data are represented as mean±SEM. p = 0.0298, two-tailed paired t-test. (G) Immunofluorescence staining of degron (red) in epidermis of K10-Degron;H2B-GFP neonates and control H2B-GFP littermate. Insets show only GFP in control (1) or in K10-Degron;H2B-GFP (2). (H) Quantification of average H2B-GFP fluorescence intensity per nucleus in controls vs degron+ cells in experimental neonates (n=49 cells for control and n=63 degron+ cells for K10-Degron;H2B-GFP from 2 animals per genotype). p < 0.0001, two-tailed unpaired t-test. (I) Immunofluorescence staining of GFP and degron in the tail epidermis of a K14-Degron;ZO1-GFP adult mouse. Insets show areas across cell-cell junctions between: (1) two adjacent cells not expressing degron (Degron^Neg^), (2) Degron^Neg^ and a cell expressing degron (Degron^Pos^), or (3) two adjacent Degron^Pos^ cells. (J) GFP fluorescence intensity in the adult tail epidermis across 10 cell-cell junctions for each category in (I). Line scans were performed as represented with white lines in insets 1, 2 and 3. (K) Quantification of fluorescence intensity at suprabasal-suprabasal cell boundaries in the adult tail epidermis. Each dot in the violin plots represents the maximum intensity across cell-cell junctions in each category shown in I (n=30 measurements across 10 cell-cell junctions per category). p < 0.0001, ordinary one-way ANOVA, Tukey’s multiple comparisons test.

To assess the ability of Degron^GFP^ to knock down GFP-tagged proteins, we began by crossing it to a mouse line with a functional knock-in of GFP in frame with ZO1 (Foote et al., 2013),a tight junction protein that localizes to the cell cortex. Further, we used a Krt10-rtTA line to direct expression of Degron^GFP^ in the differentiated layers of the epidermis (Muroyama & Lechler, 2017). An anti-camelid nanobody antibody was used to detect the expression of VHL-aGFP (hereafter referred to as degron), which was found uniformly throughout the suprabasal cells of the epidermis but was not detected in the stem/progenitor cells of the basal layer (Fig 1B,E,G). We quantified ZO1-GFP levels in neonatal K10-Degron; ZO1-GFP^Het^ mice using controls from the same litter that were degron negative. While there was a very clear cortical localization of ZO1-GFP in controls, we were unable to detect any signal in the degron positive cells of Degron^GFP^-expressing embryos(Figure 1B-D). Notably, ZO1 was degraded along with the GFP as in a homozygous ZO1-GFP mouse, endogenous ZO1 was also lost (Supplementary Figure 1).

We next asked whether Degron^GFP^ could degrade GFP-tagged proteins that were targeted to different subcellular sites. Using either a membrane-targeted GFP (mGFP) or a nuclear GFP (Krt14-H2B-GFP), we saw efficient GFP degradation in these very different compartments wherever degron was expressed (Figure 1E-H). The levels of GFP in the H2B-GFP line are by far the highest of the three, and histones are very stable proteins, thus demonstrating the efficiency of the system, even in this membrane-delimited subcellular compartment.

While degron expression was usually uniform in Degron^GFP^ embryos, we found examples in adult animals where its expression was mosaic, especially in the tail skin. This allowed us to observe that the ZO1-GFP levels in junctions between two adjacent cells expressing degron (Degron^Pos^) were drastically reduced compared to two adjacent degron-negative (Degron^Neg^) cells. GFP intensity was decreased to about half between Degron^Pos^/Degron^Neg^ junctions in which only one side of the cell junction has ZO1-GFP depleted (Figure 1I-K). This type of mosaic analysis, which can be achieved utilizing the dose-dependent inducible CreER system, could be useful for looking at mutant cells in a wild-type background.

### Degron^GFP^ degrades GFP-tagged proteins in distinct cell types

To test whether Degron^GFP^ can be useful for targeted protein degradation in other cell populations beyond the differentiated epidermis, we crossed it with different cell-type specific rtTA lines. In addition to the Krt10-rtTA line, we also tested rtTA lines under the promoters Krt14, expressed in basal progenitor cells of the epidermis, and Villin, expressed in intestinal epithelial cells. In the Krt10-rtTA model, only suprabasal cells were targeted, but GFP expression remained in basal cells (Figure 1B). Because ZO1-GFP levels are quite low in basal cells, we instead tested degradation in this layer using glucocorticoid receptor-GFP (GR-GFP) mice (Brewer et al., 2002). GR-GFP is expressed throughout the epidermis and localizes to the nucleus. Using the Krt14-rtTA line, we found expression of Degron^GFP^ specifically within the basal progenitor cells, but not in the differentiated layers. Consistent with this finding, we observed a loss of GR-GFP only in the basal layer of the epidermis. Thus, while the progenitor basal cells had depleted GR-GFP, their progeny still expressed it (Figure 2A, C). While the K14-rTA line allowed us to specifically target basal cells in the embryo, this line did not show tight restriction to this layer in the adult (Figure 1I).

**Figure 2.**
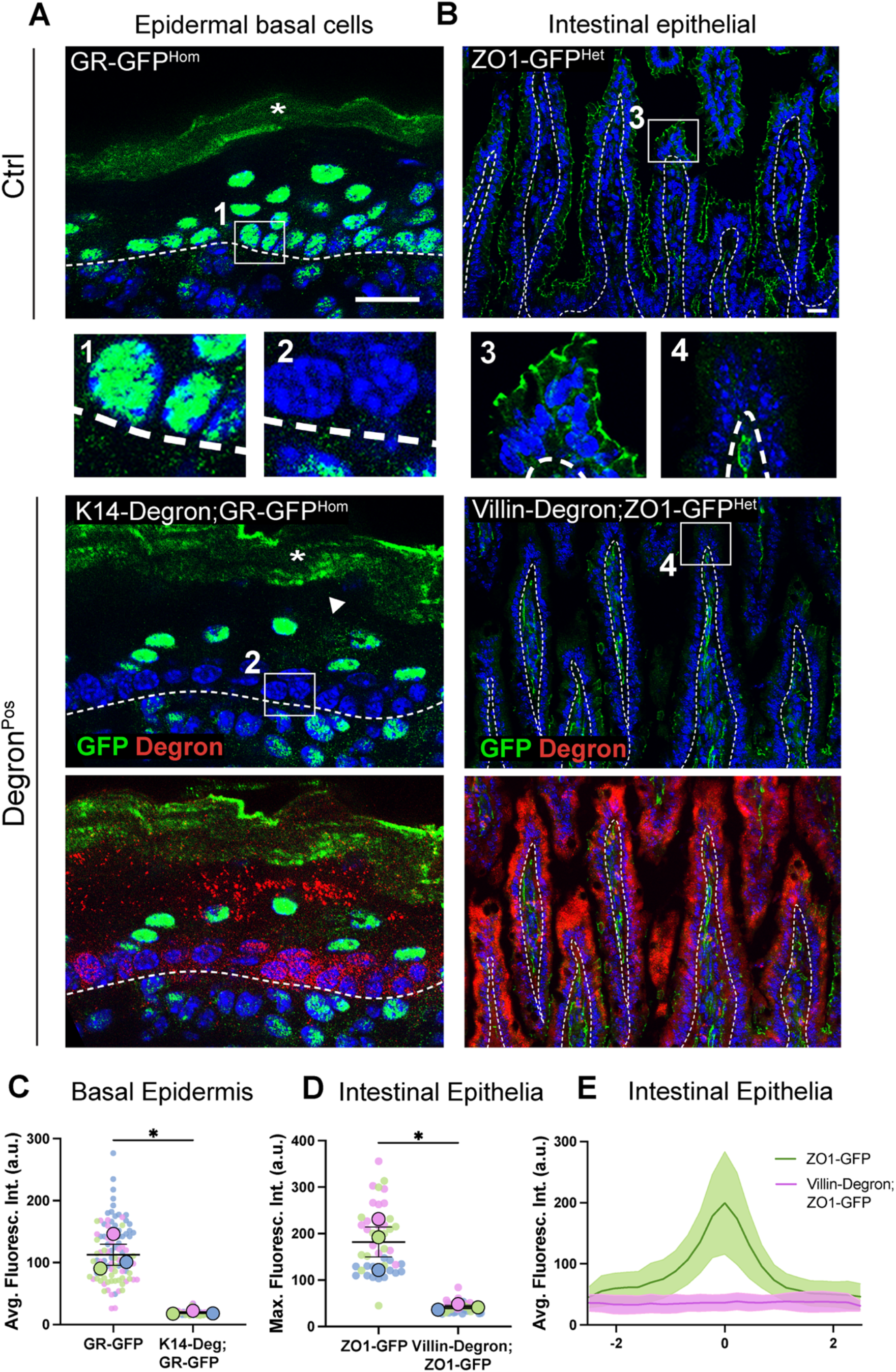
Degron^GFP^ degrades GFP-tagged proteins in different cell populations (A) Immunofluorescence staining of GFP (green) and degron (red) in E18.5 back skin epidermis of K14-Degron;GR-GFP and control GR-GFP littermate. Insets show only GFP in control (1) or in K10-Degron;ZO1-GFP (2). White arrowhead indicates unspecific antibody signal. Dotted lines represent the basement membrane. *: Indicates auto-fluorescent cornified layer. Scale: 20μm. (B) Immunofluorescence staining of GFP and degron in adult intestinal epithelia of Villin-Deg; ZO1-GFP and control ZO1-GFP littermate. Scale: 20μm. (C) Quantification of nuclei average fluorescence intensity in the basal layers in GR-GFP (n=3 mice, 98 nuclei total), or degron^Pos^ basal cells in K14-Degron;GR-GFP (n=3 mice, 110 nuclei total) embryos. Dots represent each nucleus measured, large circles represent the average for each mouse. Lines show the mean±SEM. p=0.0268, paired two-tailed t-test. (D) Quantification of fluorescence intensity across apical cell-cell junctions in villi epithelial of ZO1-GFP, and degron^Pos^ areas in Villin-Degron;ZO1-GFP neonates. Each dot represents the maximum intensity at cell-cell junctions, large circles represent the average for each mouse (n=3 mice per genotype, 15 line scans per animal). Lines show the mean±SEM. p=0.0294, paired two-tailed t-test. (E) GFP fluorescence intensity across cell-cell villi epithelial cells of ZO1-GFP (green) and degron^Pos^ areas of Villin-Degron;ZO1-GFP mice (pink). Line scan graph shows measurements across 12 cell-cell boundaries from 3 different mice per genotype. Data are represented as mean±SD.

We also tested this mouse line in the intestine, an epithelial tissue with a very high turnover rate. Using a Villin-rtTA line (Gao et al., 2013), we induced degron expression in villar epithelial cells of the intestine, although we occasionally observed it in the crypts as well (Fig 2B). ZO1-GFP, which marks tight junctions at the apicolateral membrane of enterocytes, was significantly decreased in regions of the intestine where degron was present in Villin-Degron; ZO1-GFP^Het^ mice (Figure 2B, D, E). Thus, Degron^GFP^ can deplete GFP-tagged proteins in different tissues.

### GFP-tagged protein degradation is relatively rapid and reversible

One advantage of the degron systems is the rapid loss of protein. However, the doxycycline-inducible system has an initial requirement for transcription and translation of the degron, which makes it slower than more direct pharmacologically-induced degradation. To assess the kinetics of Degron^GFP^-based degradation *in vivo*, we treated mice with different durations of doxycycline exposure. When K10-Degron;ZO1-GFP^Het^ embryos were exposed to dox for 1 day, we observed complete loss of detectable GFP signal (Figure 3A,C). To challenge the system, we also examined GFP levels 6 hours after doxycycline treatment. At this early time point, degron expression was much lower and more difficult to detect. However, the low levels resulted in loss of GFP signal, similar to doxycyline-exposure for 24 hours and negative control WT samples (Figure 3B, D). These data demonstrate that Degron^GFP^ can efficiently deplete GFP-fused proteins in as little as 6 hours after induction *in vivo*.

**Figure 3.**
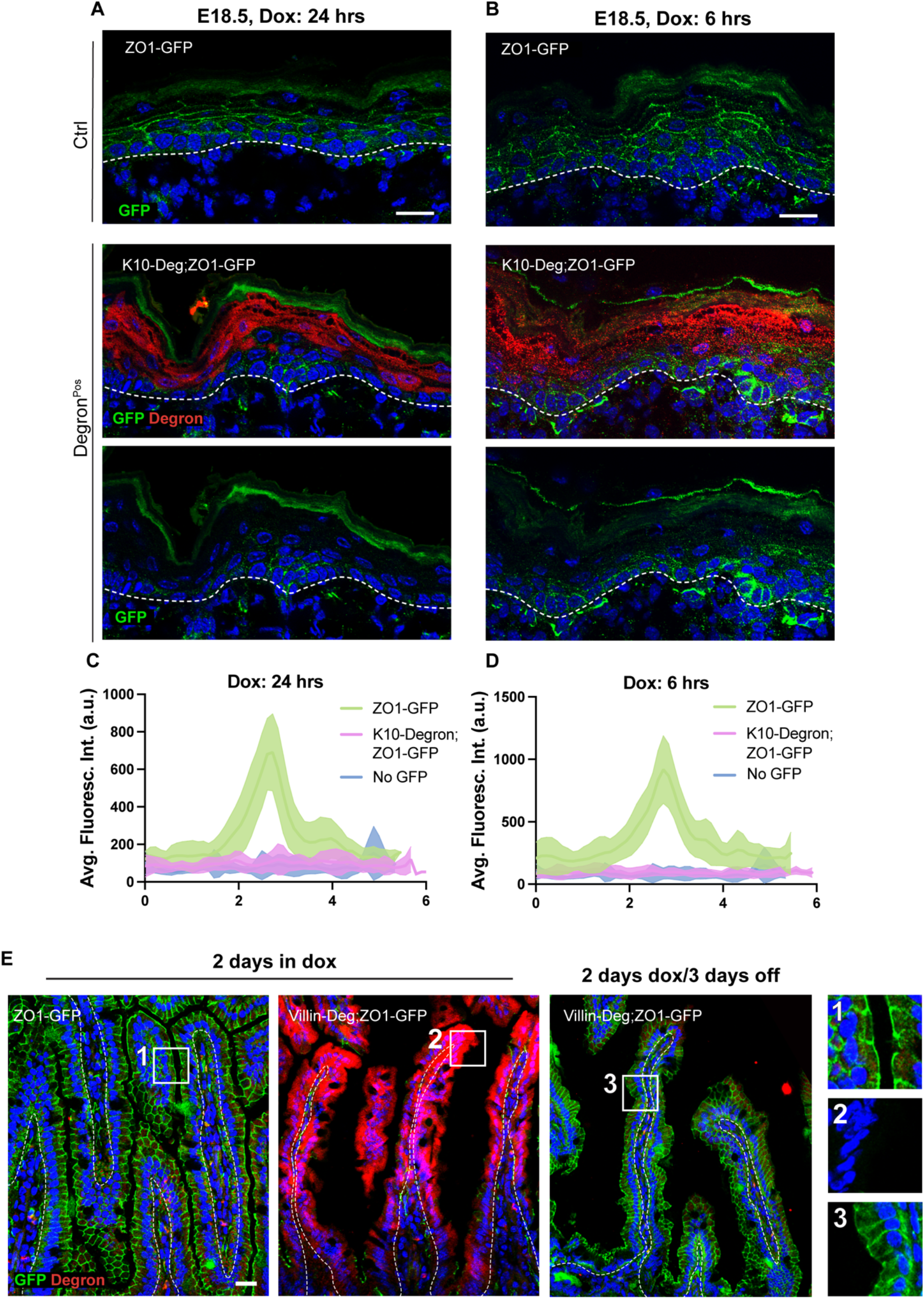
Kinetics of GFP degradation using Degron^GFP^. (A and B) Immunofluorescence staining of GFP (green) and degron (red) in E18.5 back skin epidermis of K10-Degron;ZO1-GFP and control ZO1-GFP embryos 24 hours (A) or 6 hours (B) after dox exposure. Dotted lines represent the basement membrane. All scales: 20μm. (C and D) GFP fluorescence intensity across cell-cell junctions in the epidermal suprabasal layers of ZO1-GFP (green), suprabasal degron^Pos^ areas of K10-Degron;ZO1-GFP (pink) neonates 24 hours (C) or 6 hours (D) after dox exposure, and negative control WT (blue). Line scan graph shows measurements across nine cell-cell boundaries from 3 different mice per genotype. Data are represented as mean±SD. (E, F and G) Immunofluorescence staining of GFP (green) and degron (red) in adult intestinal epithelia of 2 days dox-exposed Villin-Deg; ZO1-GFP (E), control ZO1-GFP (F), and Villin-Deg; ZO1-GFP out of dox for 3 days (G). Scale: 20μm. E) Immunofluorescence staining of GFP (green) and degron (red) in adult intestinal epithelia after 2 days of doxycycline in control ZO1-GFP (left panel, inset 1) and Villin-Deg;ZO1-GFP (middle panel, inset 2). Right panel, inset 3, shows Villin-Deg;ZO1-GFP that was taken off of doxycycline for 3 days. Insets show only GFP. Scale: 20μm.

To test the reversibility of this system, we evaluated whether GFP expression returned after stopping doxycycline exposure. For this, we used Villin-Degron; ZO1-GFP^Het^ mice, which after 3 days without doxycycline exhibited GFP signals at similar levels to their control littermates (Fig 3E,F). Kinetics of return will likely be highly variable between cell types and proteins, as they are dependent on degron stability, tissue turnover, and rate of production of new protein.

### Degron^GFP^ recapitulates GR knockout phenotypes and identifies cell types where it functions

Next, we sought to assess whether Degron^GFP^ would be useful for performing loss of function studies *in vivo*. We chose to test this on the Glucocorticoid receptor (GR) because: 1. it is expressed throughout all layers of the epidermis, 2. it has reported developmental phenotypes in both basal and differentiated cells in the context of knockout models, and 3. there was an existing GFP-knock in line. GR is part of the nuclear receptor superfamily, and upon binding to its ligand (including glucocorticoids), it acts as a transcription factor (Perez, 2011).

Previous studies of null and epidermis-specific GR knockouts (GR^eKO^) have reported epidermal differentiation defects during development, as well as aberrant proliferation (Bayo et al., 2008; Latorre et al., 2013; Sevilla et al., 2010; Sevilla et al., 2013). These reports demonstrate multiple roles of GR in the epidermis. However, the specific role of GR in each cellular compartment of the epidermis and how they contribute to the aforementioned effects still remained unanswered. We began by asking whether we were able to specifically remove GR from basal verses suprabasal cells.

We generated homozygous GR-GFP mice that also expressed degron under either a basal(K14) or suprabasal (K10) promoter. When we used K14-Degron;GR-GFP^Hom^ to induce targeted GR degradation, we found a specific loss of signal within the basal cells, but suprabasal cells maintained their GFP signal (Figure 4B, G). The reverse was true using the K10-Degron system – basal GR was maintained, while signal in differentiated cells was lost (Figure 3C, H). GR was degraded along with GFP as antibodies to endogenous GR revealed no signal in the presence of Degron^GFP^ (Supplementary Figure 1).

**Figure 4.**
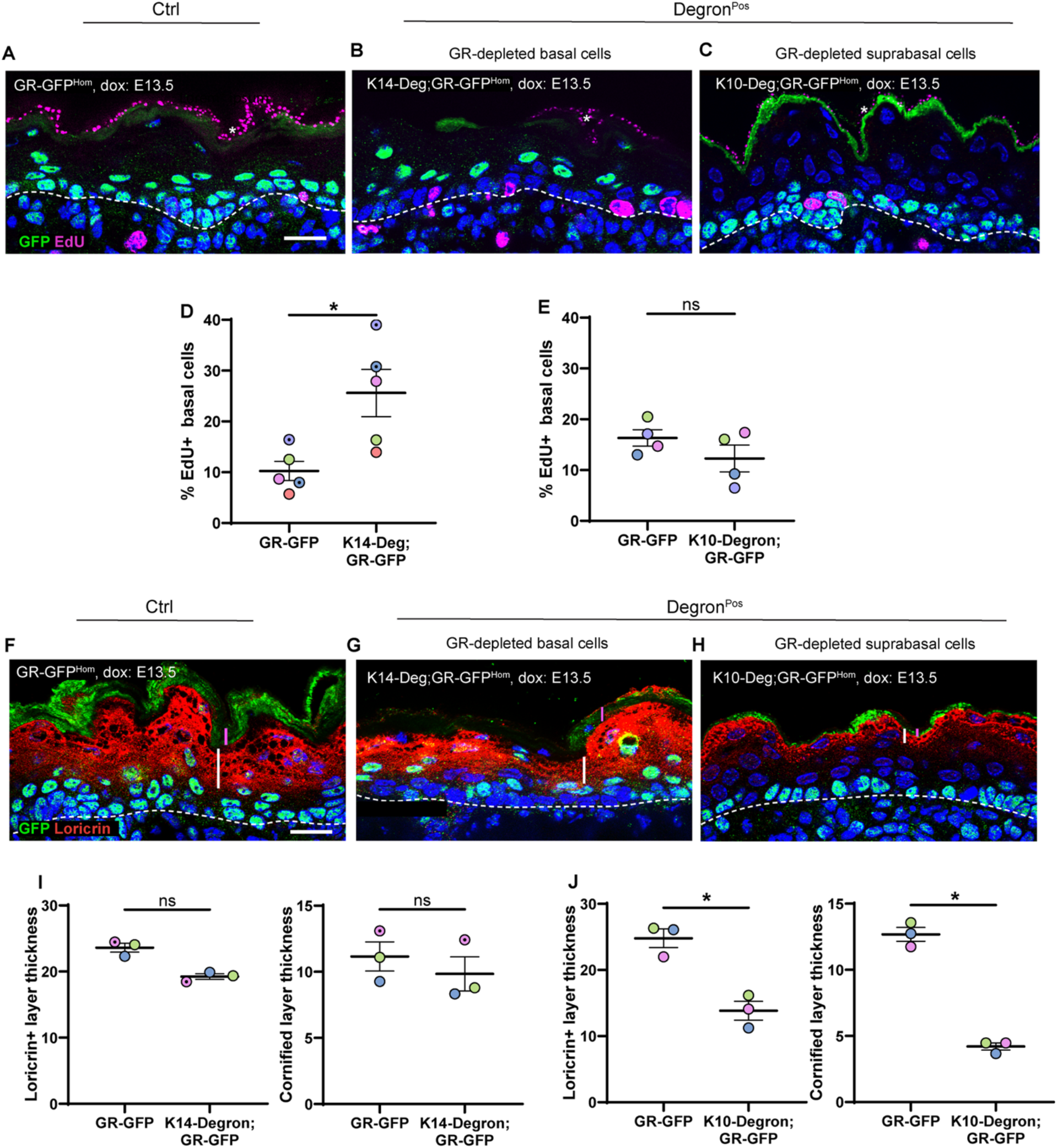
Degron^GFP^ efficiently depletes GR-GFP and reproduces knockout phenotypes. (A,B,C) Immunofluorescence staining of GFP (green) and EdU (magenta) in E18.5 back skin epidermis of GR-GFP (A), K14-Degron;GR-GFP (B) and K10-Degron;GR-GFP (C) embryos. *: Indicates non-specific signal in cornified layer. Scale: 20μm. (D) Percentage of Edu positive cells in K14-Degron;GR-GFP^Hom^ vs GR-GFP control littermates. Each circle represents measurements of a mouse: solid color circles represent mice dox-exposed since E13.5, and circles with a dot represent dox-exposed mice since E16.5 (n=5 mice per genotype). Lines show the mean±SEM. *: p=0.0172, paired two-tailed t-tests. (E) Percentage of Edu positive cells in K10-Degron;GR-GFP^Hom^ vs GR-GFP control littermates. Each circle represents measurements of a dox-exposed mouse since E13.5 (n=4 mice per genotype). Lines show the mean±SEM. *: p=0.0235, paired two-tailed t-test. (F,G,H) Immunofluorescence staining of GFP (green) and Loricrin (red) in E18.5 back skin epidermis of GR-GFP (F), K14-Degron;GR-GFP (G) and K10-Degron;GR-GFP (H) embryos. White and pink lines show granular and cornified layers thickness, respectively. Dotted white lines represent the basement membrane. Scale: 20μm. (I) Quantification of loricrin and cornified layer thickness in GR-GFP controls vs K14-Degron;GR-GFP^Hom^ littermates. Each circle represents the average layer thickness of a mouse: solid color circles represent mice dox-exposed since E13.5, and circles with a dot represent dox-exposed mice since E16.5. Lines show the mean±SEM. Loricrin and cornified thickness measurements were taken perpendicular to the basement membrane (n=3 mice per genotype, at least 22 measurements per mouse) p=0.0528 and 0.1206, respectively. ns: non-significant, paired two-tailed t-tests. (J) Quantification of loricrin and cornified thickness in GR-GFP controls vs K10-Degron;GR-GFP^Hom^ littermates. Each circle represents the average loricrin or cornified thickness of a mouse dox-exposed since E13.5. Lines show the mean±SEM (n=3 mice per genotype, at least 30 measurements per mouse) *: p=0.033 and 0.1206. ns: non-significant, paired two-tailed t-tests.

Having established the lines, we next asked about the phenotypic consequences of loss of function in each of these distinct cellular compartments. We first assessed basal cell proliferation as epidermal knockout of GR^eKO^ resulted in hyperproliferation (Sevilla et al., 2013). There was a significant increase in basal cell proliferation in the K14-Degron GR mice but not in the K10-Degron strain (Fig 4A-E). This demonstrates that the effects on proliferation are cell-autonomous and not due to defective signaling from daughters or disrupted tissue function.

We next examined differentiation by staining for the granular differentiation marker, loricirin, whose expression is defective in the GR^eKO^ mice (Sevilla et al., 2013). In contrast to proliferation, loss of GR in basal cells had no effect on differentiation as assessed by either the thickness of the granular (loricrin positive) layer, or of the cornified envelope (Fig 4F-J). In this case, however, there was a clear role for suprabasal GR, as there was a significant decrease in both granular and cornified thickness in the K10-Degron;GR-GFP^Hom^ mice (Bayo et al., 2008; Sevilla et al., 2010; Sevilla et al., 2013). Overall, knockdown studies using Degron^GFP^ were able to replicate phenotypes observed in traditional knockouts. For the first time, this tool allowed loss of function analysis of GR in different compartments of the epidermis, demonstrating distinct roles in different cell types contributing to the pleiotropic effects seen in GR knockouts. Importantly, these data show that Degron^GFP^ is a useful tool for loss-of-function studies *in vivo*.

## Discussion

We have developed a mouse line that allows for degradation of GFP-tagged proteins.

This system is rapid, reversible, and allows for both temporal and spatial control of protein degradation.

In comparison to recombination-based systems, protein degradation is more rapid and is reversible. Existing degron alleles in mouse (including AID-based and dTAG) offer faster protein degradation than the Degron^GFP^ described here. However, Degron^GFP^ offers the advantage of controlling degradation in a cell-type specific manner, alongside the timing of degradation. In addition, the AID and dTAG require a specific tag with a gene of interest, which entails the creation of a new mouse line that is monofunctional for degradation. In contrast, this system uses GFP as a tag for targeted degradation, which can also enable the examination of protein expression, localization and dynamics. Additionally, it takes advantage of existing GFP knock-in lines. Moreover, the inducer molecule, doxycycline, is well tolerated in mice and crosses the blood/brain barrier. It thus offers some advantages to the pharmacological activators of other degrons (Macdonald et al., 2022; Suski et al., 2022; Yesbolatova et al., 2020). Although a notable limitation of the Degron^GFP^ is that it requires two transgenes (rtTA and Degron^GFP^), instead of the single transgene required for degradation in other systems, Degron^GFP^ allows for tissue specificity and reversibility.

In terms of functional validation, we have used Degron^GFP^ to knockdown GR expression in two different cell compartments within the skin epidermis. Because GR is expressed throughout this tissue, previous knockout approaches could not identify in which cell types GR activity was required. Using Degron^GFP^, we were able to deplete a protein in the stem cell pool for the first time without altering its expression in the differentiated cells progeny. This demonstrated that GR has distinctive roles in basal and differentiated cells to locally control both proliferation and differentiation. This work demonstrates that Degron^GFP^-induced knockdown can recapitulate existing knockout phenotypes and provide more flexibility in defining a gene’s role in different cell types and times of activity. It is both time-consuming and expensive to generate new mouse lines, which is one reason why there are limited GFP knock-in lines for analysis, despite their great utility in understanding cell and tissue biology. The ability to use GFP knock-in lines for loss of function analysis may provide additional motivation for generation of these useful tools.

## Materials and Methods

### Mice

All mouse work was approved by Duke University’s Institutional Animal Care and Use Committee and performed in accordance with committee guidelines. Mice were genotyped by PCR, and both male and female mice were analyzed in this study. All mice were maintained in a barrier facility with 12-hour light/dark cycles. Mouse strains used in this study include: K10-rtTA (Muroyama & Lechler, 2017), ZO1-GFP (Foote et al., 2013), K14-H2BGFP (Tumbar et al., 2004), GR-GFP (from Dr. Diana Laird, (Brewer et al., 2002)). Other mice were from the Jackson Laboratories, and their stock numbers are as follows: K14-rtTA (008099), Villin-rtTA (031283/ 031285), mGFP was generated by mating Rosa-mT/mG (007576) mice with a CMV-Cre-deleter strain (006054) (Ning et al., 2021).

### Generation of GFP^Degron^ mouse line

To generate the Degron^GFP^ line, we used the pBABED Hygro Tet-on VHL aGFP plasmid (MRC PPU, DU 54221). To verify the proper doxycycline-dependent expression of this plasmid, the vector was co-transfected with a K14-rtTA plasmid (Ning et al., 2021) into cultured keratinocytes and placed in doxycycline-containing media. The plasmid was linearized, purified, and used by the Duke Transgenic Core to generate transgenic mice via pronuclear injection.

Genotyping to identify founders was performed using forward primer: CTGCCCGTATGGCTCAACTTC, and reverse primer: CTAGAAACCGTCACCTGGG. Two lines were kept which have robust transgene expression in the intestine and the epidermis, respectively.

### Immunofluorescence staining

Fresh tissue was embedded in OCT (Sakura), frozen, and sectioned at 10 μm using a Cryostat. Sections were fixed with 4% paraformaldehyde (PFA) in PBS for 8 minutes at room temperature, washed with PBS containing 0.2% Triton (PBST) for 5 minutes, then blocked with blocking buffer (3% bovine serum albumin with 5% normal goat serum (Gibco,16210064), and 5% normal donkey serum (Sigma-Aldrich, D9663) in PBST) for 15minutes. Sections were incubated with primary antibodies diluted in blocking buffer overnight at 4°C (or at room temperature: 1hr for Loricrin, 2hrs for GR), then three times washed with PBST, and incubated in secondary antibodies and Hoechstv34580 for 15 minutes at room temperature. After three washes with PBST, sections were mounted in the anti-fade buffer (90% glycerol in PBS plus 2.5 mg/ml p-Phenylenediamine (Thermo Fisher, 417481000) and sealed using clear nail polish in the borders.

Primary antibodies used in this study: mAb Rb anti VHH-iFluor555 (1:500, GenScript, A01863), Ch anti GFP (1:2000, Abcam, ab13970), Rb anti Loricrin (1:100, gift from Colin Jamora), mAb Rb anti GR (1:100, Cell Signaling, 12041T).

### EdU labeling

Pregnant dams were intraperitoneally injected with 10 mg/kg of EdU and sacrificed after 1 hour for tissue dissection. Back skins of embryos were collected and fresh frozen, and tails were taken for genotyping. Tissue sections were fixed with 4% PFA and stained with primary and secondary antibodies, then EdU was detected following Click-iT EdU (Thermo Fisher,C10337) protocol.

### Granular and cornified measurements

To measure granular thickness, perpendicular lines to the basement membrane were traced along the Loricrin positive area in 15 different spots across each field of view using the straight-line tool in Fiji and measured their length. Similarly, 15 lines per field of view were traced across background-staining of the cornified layer in the green channel to measure cornified layer thickness.

---

**Supplementary Figure 1.**
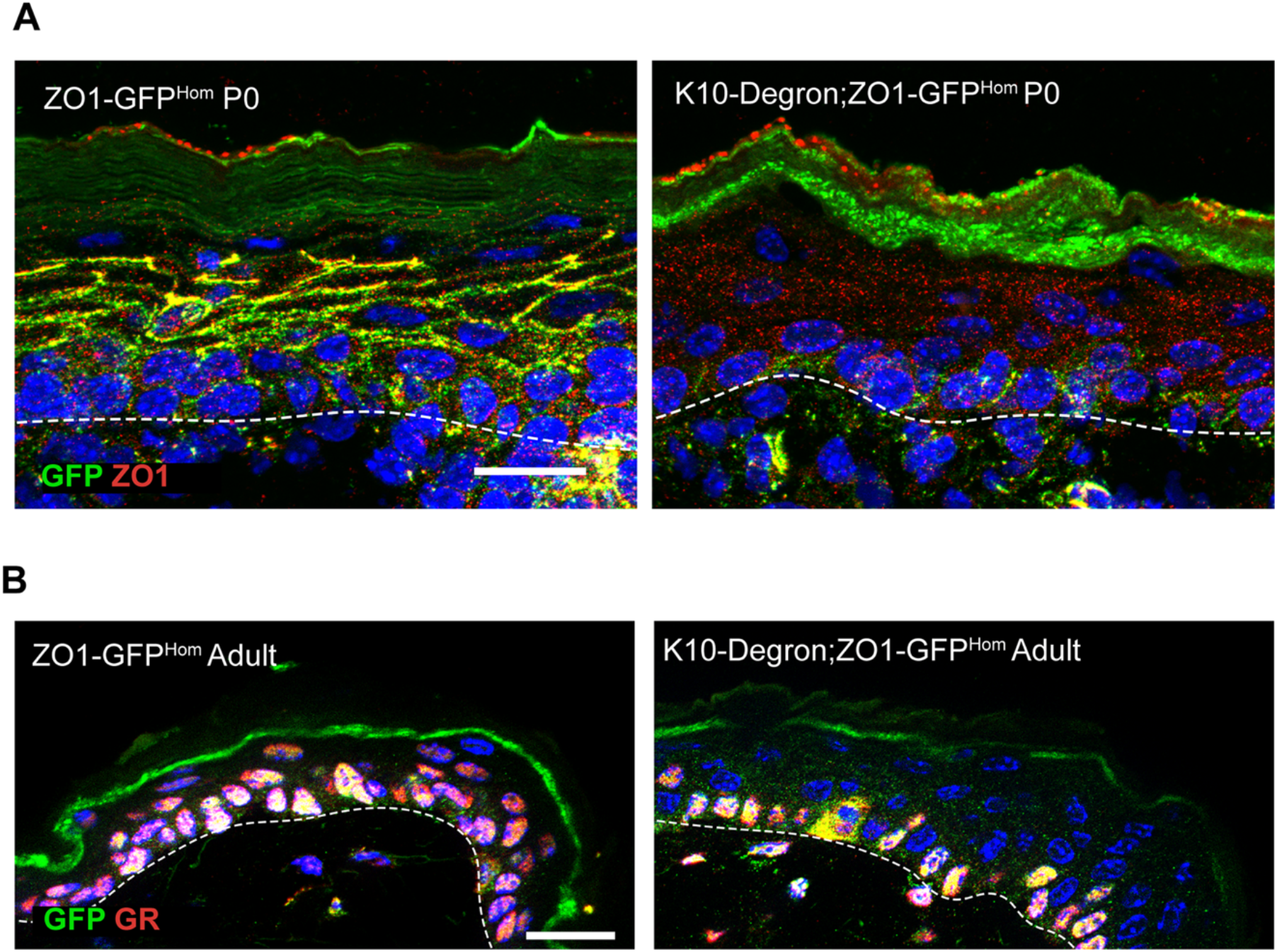
Degron^GFP^ degrades both GFP and the protein fused to it. (A) Immunofluorescence staining of GFP (green) and ZO1 (red) in P0 back skin epidermis of K10-Degron;ZO1-GFP and control ZO1-GFP neonates. Dotted lines represent the basement membrane. Scale: 20μm. (B) Immunofluorescence staining of GFP (green) and GR (red) in adult back skin epidermis of K10-Degron;GR-GFP and control ZO1-GFP littermate. Dotted lines represent the basement membrane. Scale: 20μm.

